# AATF Inhibition Exerts Antiangiogenic Effects Against Human Hepatocellular Carcinoma

**DOI:** 10.1101/2022.11.28.518287

**Authors:** Diwakar Suresh, Akshatha N. Srinivas, Akila Prashant, Suchitha Satish, Prashant Vishwanath, Prasanna K. Santhekadur, Divya P. Kumar

**Affiliations:** Department of Biochemistry, CEMR, JSS Medical College, JSS Academy of Higher Education and Research, Mysuru, Karnataka, India; Department of Pathology, JSS Medical College and Hospital, JSS Academy of Higher Education and Research, Mysuru, India

**Keywords:** Apoptosis antagonizing transcription factor, angiogenesis, hepatocellular carcinoma, knockdown, human umbilical vein endothelial cells, pigment epithelium-derived factor

## Abstract

Hepatocellular carcinoma (HCC), a highly fatal cancer with a mortality rate proportional to its incidence, continues to pose a global health care challenge. Angiogenesis is a key factor in the growth and metastasis of hepatic tumors and thus a potential therapeutic target in HCC. We previously showed that the apoptosis antagonizing transcription factor (AATF) affects tumor growth and metastasis in a mouse xenograft model. However, the regulatory role of AATF in tumor angiogenesis and its underlying mechanisms in HCC remain unknown. In the current study, we identified high levels of AATF in human HCC tissues compared to adjacent normal liver tissues, and the expression was found to be correlated with the stages and tumor grades of HCC. Inhibition of AATF in human HCC cells showed high levels of pigment epithelium-derived factor (PEDF) compared to controls as a resultant of reduced matric metalloproteinases activity. Conditioned media from AATF knockdown (KD) cells inhibited the proliferation, migration, and invasion of human umbilical vein endothelial cells as well as the vascularization of the chick chorioallantoic membrane. Furthermore, the VEGF-mediated downstream signaling pathway responsible for endothelial cell survival and vascular permeability, cell proliferation, and migration favoring angiogenesis was suppressed by AATF inhibition. Of note, PEDF inhibition by its respective antibody profoundly reversed the anti-angiogenic effect of AATF KD. In conclusion, our study demonstrates that inhibition of AATF suppresses angiogenesis in HCC via PEDF. Thus, a therapeutic strategy based on the inhibition of AATF to disrupt tumor angiogenesis may be a promising approach for HCC treatment.

## INTRODUCTION

Hepatocellular carcinoma (HCC), which makes up 80% of primary liver malignancies, has become a severe public health issue. It is estimated that in the next two decades, there will be a 55% increase in the incidence rate of liver cancer, posing a challenge worldwide (1). The underlying causes of HCC development have been identified as viral infections (Hepatitis B virus and Hepatitis C virus), as well as additional risk factors including metabolic syndrome, carcinogens, fatty liver disease, and cirrhosis (2). Hepatic injury and chronic liver inflammation cause hepatocyte necrosis, regeneration, and the progression to fibrosis, cirrhosis, resulting in the onset and progression of HCC (3). The pathophysiology of HCC is multifactorial and highly complex owing to its molecular and immune heterogeneity, and thus, understanding the molecular processes could facilitate the development of preventive measures, early diagnostic techniques, and improved therapeutic options (4,5). HCC being a highly vascular tumor underscores the importance of angiogenesis in the process of tumor growth and metastasis, which is responsible for the rapid recurrence and poor survival rates of HCC (6).

Angiogenesis is one of the hallmarks of cancer and has a crucial role in the development of solid malignancies (7). The process of angiogenesis aids cancer cells in creating a local vascular ecology to deliver nutrients and growth factors, remove potentially toxic metabolites, and thereby promote cancer cell proliferation and metastasis (6). Angiogenesis is stimulated by various proangiogenic factors such as vascular endothelial growth factor (VEGF), platelet-derived growth factor (PDGF), fibroblast growth factor (FGF), and angiopoietins (8). The molecular understanding of this intricate and dynamic angiogenic tumor ecosystem has led to the advancement of anti-angiogenic therapy for HCC (9). The “starve a tumor to death” theory has emerged as an appealing anti-angiogenic therapy for a variety of cancers, including HCC (10). Sorafenib, a multikinase inhibitor, exerts an anti-tumor effect by inhibiting angiogenesis. In addition, over the last decade, ramucirumab and bevacizumab have been the FDA-approved drugs that target vascular endothelial growth factors (VEGFs) to treat HCC (11,12,13). Thus, the strategies targeting angiogenesis have been considered effective in treating HCC development and progression.

Apoptosis antagonizing transcription factor (AATF), also called Che-1, is a transcription factor that controls several genes involved in the regulation of different processes such as cell proliferation, cell cycle arrest, DNA damage response, and apoptosis (14). Previous studies have shown that AATF regulates the transcription of many genes, including nuclear hormone receptor-targeted genes, p53, p21, and the X-linked inhibitor of apoptosis (15–18). Additionally, AATF also plays a role in the pathogenesis of many cancers. AATF levels were found to be elevated in various cancers such as breast cancer, leukemia, lung cancer, Wilm’s tumor, osteosarcoma, and neuroblastoma (19–24), while AATF levels were downregulated in colon cancer. AATF is well studied as a component of the unfolded protein response (UPR), which is an adaptive mechanism activated during endoplasmic reticulum (ER) stress. AATF induced as a resultant of ER stress protects the cells from apoptosis by activating the transcription factor Akt1 (25). An interesting study by Wang et al., has shown that AATF alleviates hypoxia/reoxygenation (H/R)-induced cardiomyocyte apoptosis by upregulating Nrf2 signaling (26). Shimizu et al., recently reported that elevated NRAGE expression is significantly correlated with AATF expression in accelerating cancer proliferation and migration, leading to hepatocarcinogenesis (27). However, the potential role of AATF in HCC pathogenesis has not been investigated.

We have previously, for the first time, unraveled the role of AATF as a potential driver of HCC in NAFLD and demonstrated that the knockdown of AATF inhibited tumor growth and metastasis (28). In the present study, we investigated whether suppression of AATF expression inhibits angiogenesis in HCC and explored its underlying mechanisms. The specific objectives of the study were to (i) confirm the overexpression of AATF in human HCC tissues and correlate AATF expression with different stages and grades of HCC; (ii) define the impact of AATF knockdown on key angiogenic properties of HCC; and (iii) identify the signaling pathway by which AATF inhibition suppresses angiogenesis in HCC. In this study, we showed the overexpression of AATF in human HCC tissues and evaluated the role of AATF in proliferation, migration, invasion of HUVECs, and vascular growth in a chicken embryo by testing the effect of conditioned media from control and AATF knockdown HCC cells. PEDF antibody was used to investigate the effect of PEDF on AATF-mediated angiogenesis in HCC. Our findings demonstrated that AATF inhibition exerts an anti-angiogenic effect in HCC via PEDF, and that AATF merits further investigation as a potential therapeutic target, leading to a better understanding of anti-angiogenic strategies for the treatment of HCC.

## MATERIALS AND METHODS

### Reagents

Endothelial cell growth medium (EGM), extracellular matrix gel (ECM), TRIzol, RIPA buffer, collagenase A, and AATF antibody were procured from Sigma-Aldrich (St. Louis, USA). Dulbecco’s modified eagle’s medium (DMEM), giemsa stain, hematoxylin, 8μm-transwell inserts, and 24-well plates were procured from HiMedia laboratories (India). Lipofectamine 3000, penicillin/streptomycin, glutamine, FBS were from Invitrogen (USA); antibodies to IgG, PEDF, pAkt, pErk1/2, pFAK, Akt, Erk, and control and AATF shRNA from Santa Cruz Biotechnology, Inc. (CA, USA); antibodies to β-actin from Cell Signaling Technologies (USA); verso cDNA synthesis kit, and DyNamo Colorflash SYBR green kit from Thermo Fisher Scientific (USA); western bright ECL HRP substrate from Advansta (USA); western blotting materials were from BioRad; primers from Integrated DNA Technologies (IDT) (Iowa, USA), and all other reagents were obtained from Thermo Fisher Scientific or Sigma.

### Subjects and sample collection

Human HCC tissues (n=50) and adjacent normal tissues (n=15) were procured from the National Tumor Tissue Repository (NTTR), Tata Memorial Hospital, Mumbai, India. The study was approved by the institutional ethics committees of JSS Medical College, JSS AHER, Mysore, Karnataka, India (JSSMC/IEC/090721/28NCT/2021-22), and Tata Memorial Hospital. Clinical characteristics and biochemical parameters were evaluated. The demographic and clinicopathological data of the HCC subjects are described in **supplementary table 1**.

The human umbilical cord was procured from Shree Devi Nursing Home, Mysore. Informed consent was obtained from the subjects for the use of cells in clinical research. The study was approved by the institutional ethics committee at JSS Medical College, JSS AHER, Mysore, Karnataka, India. (JSSMC/IEC/260822/37NCT/2022-23).

### Isolation and culture of Human umbilical vein endothelial cells (HUVECs)

The freshly collected human umbilical cord (40-60 cm long, tied at both ends) from the maternity hospital was washed using 0.09% saline. After confirming the cord is devoid of hematic and physical damage, it is incubated with 2 mg/ml collagenase A solution at 37^0^C. The digested cells were collected into a tube and centrifuged at 750 g for 10 min at 4^0^C. The cells were resuspended with endothelial cell growth medium and cultured by incubating at 37^0^C in 5% CO_2_. All procedure was carried out in the cell culture hood with laminar airflow under aseptic condition. HUVECs were used from passages 3-5 to ensure their endothelial characteristics and were freshly isolated to perform experiments in duplicate or triplicate (**Supplementary Figure 1**).

### Cell culture and stable clones preparation

Human HCC cells-QGY-7703 and Hep3B were cultured in Dulbecco’s modified eagle’s medium containing 4.5 g/L glucose and supplemented with 10% fetal bovine serum, L-glutamine, and 100 U/ml penicillin-streptomycin incubated at 37^0^C in 5% CO_2_.

Stable clones expressing AATF shRNA in QGY-7703 cells were prepared as described previously (28). Firstly, the optimal antibiotic concentration for selecting the stable cell colonies was determined using different concentrations of puromycin (1-10 μg/ml). Control and AATF shRNA plasmid containing puromycin resistance gene was transfected to QGY-7703 cells according to the manufacturer’s protocol. Individual colonies were isolated, expanded and maintained in 1 μg/ml Puromycin. Stable knockdown of AATF in QGY-7703 cells was confirmed by qRT-PCR and western blot.

### Tissue processing and histological analysis

Human normal and HCC tissues were fixed in a 4% (v/v) formaldehyde solution in phosphate buffered saline for 16 h. After formalin fixation, tissues were processed and embedded in standard paraffin blocks. Subsequently, tissue sections of 5 μm thickness were cut from each paraffin block and stained with hematoxylin and eosin (H&E). A pathologist at JSS Hospital performed the histological grading of HCC according to the World Health Organization (WHO) classification: well differentiated, moderately differentiated, or poorly differentiated HCC (29). The various stages of HCC were determined according to the classification criteria of the American Joint Committee on Cancer (AJCC) TNM [primary tumor features (T), presence or absence of nodal involvement (N), and distant metastasis (M) staging systems (30).

### Immunohistochemistry

The formalin-fixed, paraffin-embedded (FFPE) tissue sections were deparaffinized and rehydrated following the treatment with xylene and a series of ethanol concentrations. After antigen retrieval with citrate buffer (pH 6) at 94°C for 15 min, followed by washing with water, the sections were incubated with 3% hydrogen peroxide for 10 min. After incubating the slides with blocking buffer for 1 hr at room temperature, the slides were incubated with AATF antibody (1:100 dilution) overnight at 4°C in a humidified chamber. The signals were developed using the polyexcel HRP/DAB detection system-one step (PathnSitu, Biotechnologies) as per the manufacturer’s protocol, and the nucleus was stained using hematoxylin. All the immunohistochemistry images were taken using an Olympus BX53 microscope.

### Condition media preparation

The AATF control and knockdown clones were cultured until cells reached 80-85% confluency. The cells were washed with PBS and then incubated in serum-free DMEM medium for 24 hours. The conditioned medium was collected and centrifuged at 2500 rpm for 5 min at 4^0^C to remove the dead cells and cell debris. The conditioned media was aliquoted and stored at −80^0^C.

### Neutralizing Anti-PEDF antibody

Control and AATF knockdown QGY-7703 cells-released PEDF in conditioned media was blocked by using an anti-PEDF antibody. A non-specific antibody was used as an isotype control. The antibodies were used at a concentration of 5 μg/ml.

### Proliferation assay

The HUVECs were seeded in 96 well plates at a density of 10^4^ cells per well and cultured up to 70% confluency. The cells were treated with the conditioned media of control and AATF knockdown QGY-7703 cells for 24 h, 48 h, and 72 h. At the end of the treatment, WST-1 assay mix (1:10 dilution) was added to each well and incubated for 1 h at 37^0^C in 5% CO_2_. The absorbance was measured at 450 nm on a multi-mode plate reader (EnSpire™ Multimode Plate Reader, Perkin Elmer) according to the manufacturer’s protocol.

### Migration assay

The HUVECs were seeded at a density of 5 X10^5^ cells per well in 6 well plates and allowed to attain 70-80% confluency. A scratch was made using a 1 ml pipette tip across the centre of the well, and the medium was removed to get rid of the detached cells. The HUVECs were incubated with conditioned media of control and knockdown cells with or without IgG and PEDF antibodies (Santa Cruz). The migration ability of the HUVECs was evaluated by measuring the gap widths at time intervals of 0 and 24 h. Images were acquired with a Zeiss Primovert inverted microscope and analyzed for the measurement of gap distance using ImageJ software.

### Invasion assay

Transwell inserts of 8 μm pore size were coated with matrigel matrix gel and placed into a 24-well plate. The HUVECs were suspended in the serum-free endothelial cell growth medium at the density of 5X10^4^ cells and seeded into each pre-coated transwell inserts. In the lower chambers of 24 well plates were added conditioned media from control and knockdown cells with or without IgG and PEDF antibodies. Cells were incubated at 37^0^C for 24 h to analyze the invasive ability of the cells. After the incubation, the non-invasive cells in the precoated transwell inserts were removed with a sterile PBS-soaked cotton swab. The invasive cells at the bottom of the transwell inserts were fixed with paraformaldehyde and stained with Giemsa stain. Images were captured using a Zeiss Primovert inverted microscope and analyzed by comparing the number of cells that had crossed the membrane between the control and knockdown conditioned media groups.

### Chick Chorioallantoic Membrane (CAM) assay

Fertilized chicken eggs were sterilized and pre-incubated at 37^0^C in 85% humidity. After 7 days of incubation, a small hole was made on the broad side of the shell, and carefully, a window of 1 cm^2^ was created. The embryos were treated with the conditioned media of control and knockdown cells. The window was sealed, and the eggs were incubated for 48 h in the humidified incubator at 37^0^C. Images were photographed using a Nikon digital camera, and angiogenesis is quantified by comparing the number of blood vessels between the control and knockdown groups.

### Enzyme-linked Immunosorbent Assay (ELISA)

The concentration of PEDF was measured in the conditioned media collected from control and knockdown QGY-7703 cells by using ELISA kit (R&D Systems, Minnesota, USA) according to the manufacturer’s protocol.

### Zymography

The protease activity of the metalloproteases was detected by gelatin zymography. 7.5% of polyacrylamide gels containing 0.1% gelatin with a Trisglycine running buffer were used to separate proteins. After electrophoresis, gels were washed in 2.5% TritonX-100 prepared in 50 mM Tris-HCl of pH 7.5 for 1h. Later, gels were incubated in developing buffer (1% TritonX-100, 50 mM Tris-HCl pH 7.5 along with 5 mM Calcium chloride and 1 μM zinc chloride) for 24 h at 37^0^C. After the incubation, gels were stained using coomassie blue for 1 h and destained using a destaining solution. Images were taken using the Gel Doc system (Genesys) and analyzed in ImageJ software.

### RNA isolation and quantitative real-time PCR

Total RNA from the cells and frozen liver tissues was isolated using the TRIzol method. The quality and concentration of the RNA were determined by a nanodrop spectrophotometer. The RNA was reverse transcribed by following the manufacturer’s instructions using the Verso cDNA synthesis kit. The real-time PCR reactions were carried out using the DyNamo Colorflash SYBR Green kit with 0.5 mM primers (IDT), and 50 ng of cDNA in a 20 μl reaction volume. The real-time PCR reactions were performed using the Rotor-Gene Q5plex HRM System (Qiagen). The relative quantification of the mRNA fold change was calculated as 2^-ΔΔCt^ and was expressed normalized with endogenous control β-actin. The primers for the qRT-PCR were validated, and the sequences of the primers used in this study are provided in **Supplementary Table 2**.

### Immunoblotting

The lysates were prepared by homogenizing the human liver tissues and HCC cells in RIPA buffer containing protease/phosphatase inhibitors. The supernatant was collected after the homogenized tissue or cell samples were centrifuged at 13000 rpm for 10 minutes at 4^0^C. The protein concentration was determined by using the Bio-Rad protein assay dye reagent (Bio-Rad) of Bradford’s protein estimation method. A 30-50 μg of protein was loaded to separate the proteins in the SDS-PAGE and transferred onto a nitrocellulose membrane for all western blots. The membranes were blocked using 5% nonfat skim milk for an hour at room temperature and probed with specific primary antibodies (AATF, β-actin, pErk1/2, Erk1/2, pAkt, Akt, pFak, and PEDF) for overnight incubation at 4^0^C. Furthermore, membranes were washed and incubated for secondary antibodies for 2 h at room temperature. The blots were developed using the Western Bright ECL HRP substrate, and images were captured using the Uvitec Alliance Q9 chemiluminescence imaging system. The bands were quantified using ImageJ software. The intensity of each band was normalized to its respective endogenous control, β-actin.

### Gene expression analysis using TCGA database

The AATF gene expression data of normal (n= 50) and HCC (n= 371) subjects were obtained from The Cancer Genome Atlas (TCGA) and these data were then analyzed to determine whether there are differences in gene expression based on the stages and tumor grade of HCC.

### Statistical analysis

Results were calculated as means ± SEM. Statistical significance was analyzed using Student’s t-test. All statistical analyses were performed using the GraphPad Prism software (version 6), and p values < 0.05 (*, #) or < 0.001 (**, ##) were considered significant.

## RESULTS

### Upregulation of AATF in human HCC

To investigate the role of AATF in HCC, mRNA and protein levels of AATF were measured in human HCC cell lines and found to be relatively overexpressed in QGY-7703 compared to Hep3B and normal liver tissues **(Figure 1A and 1B).** Consistent with these observations, the mRNA expression of AATF in human HCC tissues (n=50) was also found to be significantly upregulated compared to adjacent normal liver tissues (n=15) **(Figure 1C)**. Histological analysis of human HCC tissues revealed distinctive changes in the cell structure and arrangement of the hepatocytes, confirming different grades of HCC **(Figure 1D)**. Furthermore, immunohistochemistry revealed that AATF expression increased gradually from stages I to IV, as well as with the differentiation grades from well differentiated to poorly differentiated HCC (**Figure E**). These findings show that AATF expression increases with the HCC stage and loss of differentiation, confirming AATF’s role in the development and progression of HCC. Next, we analyzed the publicly available TCGA database to evaluate the status of AATF in normal (n=50) and HCC (n=371). In support of our results, the data from the TCGA database provided evidence that there are significantly higher AATF expression levels in HCC compared to normal. The levels of AATF were found to be significantly related to the pathological stages and tumor grades of HCC (**Supplemental Figure 2**).

**Figure 1.**
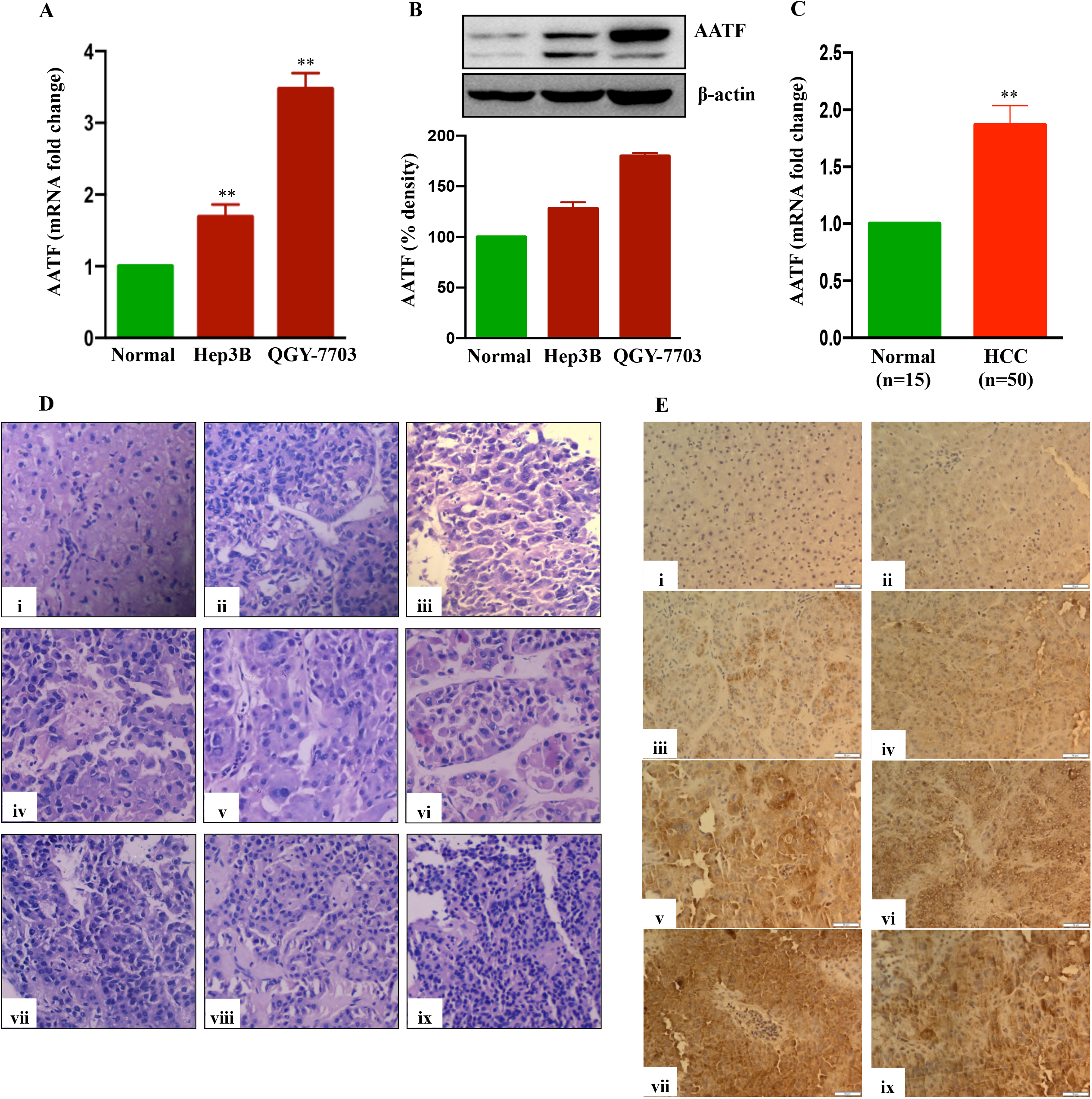
Expression of AATF in human HCC cells and HCC tissues. AATF mRNA (A) and protein (B) expression in human HCC cells. (C) AATF mRNA expression in human normal and HCC subjects. (D) Representative Hematoxylin and eosin (H&E)–stained liver sections of human normal and HCC subjects. (E) Representative immunohistochemistry images showing the expression of AATF in formalin-fixed paraffin-embedded human normal liver and HCC tissues. (i) Normal human liver; (ii) stage I, well differentiated; (iii) stage I, poorly differentiated; (iv) stage II, well differentiated; (v) stage II, poorly differentiated; (vi) stage III, well differentiated; (vii) stage III, poorly differentiated; (viii) stage IV, well differentiated; (ix) stage IV, poorly differentiated. Data are expressed as the mean ± SEM. **p < 0.001 or *p < 0.05 compared to normal.

### AATF knockdown suppresses the angiogenic potential of HCC

We investigated the regulatory role of AATF on angiogenesis, a major hallmark of cancer, which is responsible for the rapid recurrence and poor survival rate in HCC patients (31). The effect of AATF on angiogenesis was determined by assessing the impact of its loss of function. The initial step towards achieving this was the establishment of stable QGY-7703 cell lines. The expression of AATF was significantly reduced in the knockdown clones compared to puromycin-resistant control clones (**Figure 2A and Supplementary Figure 3**).

**Figure 2.**
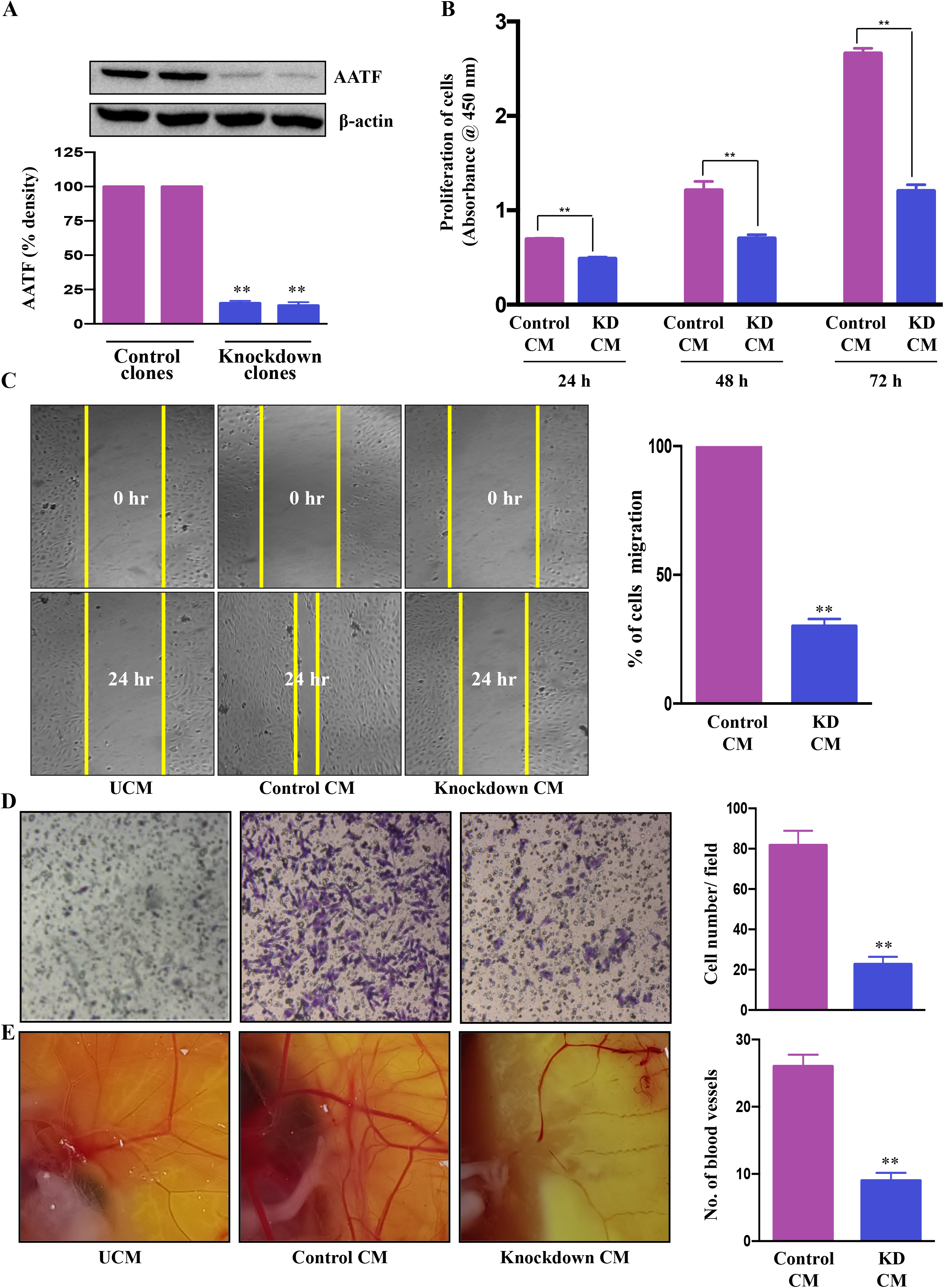
Knockdown of AATF inhibits migration, invasion of HUVECs, and vascular growth in chicken embryo chorioallantoic membrane. AATF protein (A) expression in control and AATF knockdown QGY-7703 cells. Effect of conditioned media from control and AATF knockdown QGY-7703 cells on proliferation (B), migration (C), and invasion (D) of the human umbilical vein endothelial cells (HUVECs). (E) CAM assay performed using the conditioned media from control and AATF knockdown QGY-7703 cells. Representative images are shown. UCM, unconditioned medium, CM, conditioned medium. Data are expressed as the mean ± SEM of three experiments. **p < 0.001 or *p < 0.05 compared to control.

Cell proliferation, migration and invasion are critical steps for the endothelial cells to form blood vessels in angiogenesis (32). To determine the effect of AATF on angiogenesis, the influence of conditioned media (CM) of control and AATF knockdown QGY-7703 cells on proliferation, migration, and invasion of human umbilical vein endothelial cells (HUVECs) was analyzed. We hypothesized that AATF knockdown would affect the angiogenic properties of HUVECs and vascular development in chick embryos. Consistent with this hypothesis, the CM of the knockdown cells significantly inhibited the proliferation of HUVECs compared to the control (**Figure 2B**). Furthermore, the chemotactic motility of endothelial cells was determined by migration assay and matrigel invasion assay. The results showed that CM of knockdown cells inhibited the migration of HUVECs and significantly reduced cell invasion compared to control, providing strong evidence that AATF knockdown affects the motility and matrix degradation capacity of HUVECs, which are critical for angiogenic sprouting (**Figure 2C and 2D**).

We further employed the chicken chorioallantoic membrane (CAM) assay, which is a physiological model of embryonic angiogenesis. Similar results were obtained where the CM of AATF knockdown cells showed a remarkable effect on the chicken embryo by significantly decreasing the vascular growth compared to the CM of the control cells (**Figure 2D**). Collectively, these data demonstrate that AATF knockdown exerts an antiangiogenic effect on HCC.

### Effect of AATF knockdown on PEDF levels

The proangiogenic and anti-angiogenic genes manifest themselves differently in HCC due to the activation of diverse oncogenic pathways. Our previous study using a human angiogenesis array revealed that PEDF, or SerpinF1, a well-known anti-angiogenic factor, is highly expressed in AATF knockdown cells compared to control cells (28). With this rationale, we investigated the regulatory role of AATF on PEDF levels in control and knockdown QGY-7703 stable cells. The AATF knockdown cells showed upregulated PEDF protein expression compared to controls **(Figure 3A).** Furthermore, PEDF levels were also measured by ELISA using the CM from control and knockdown cells. Similar results were obtained where the secretion of PEDF was significantly higher in the CM of AATF knockdown cells compared to the control **(Figure 3B)**.

**Figure 3.**
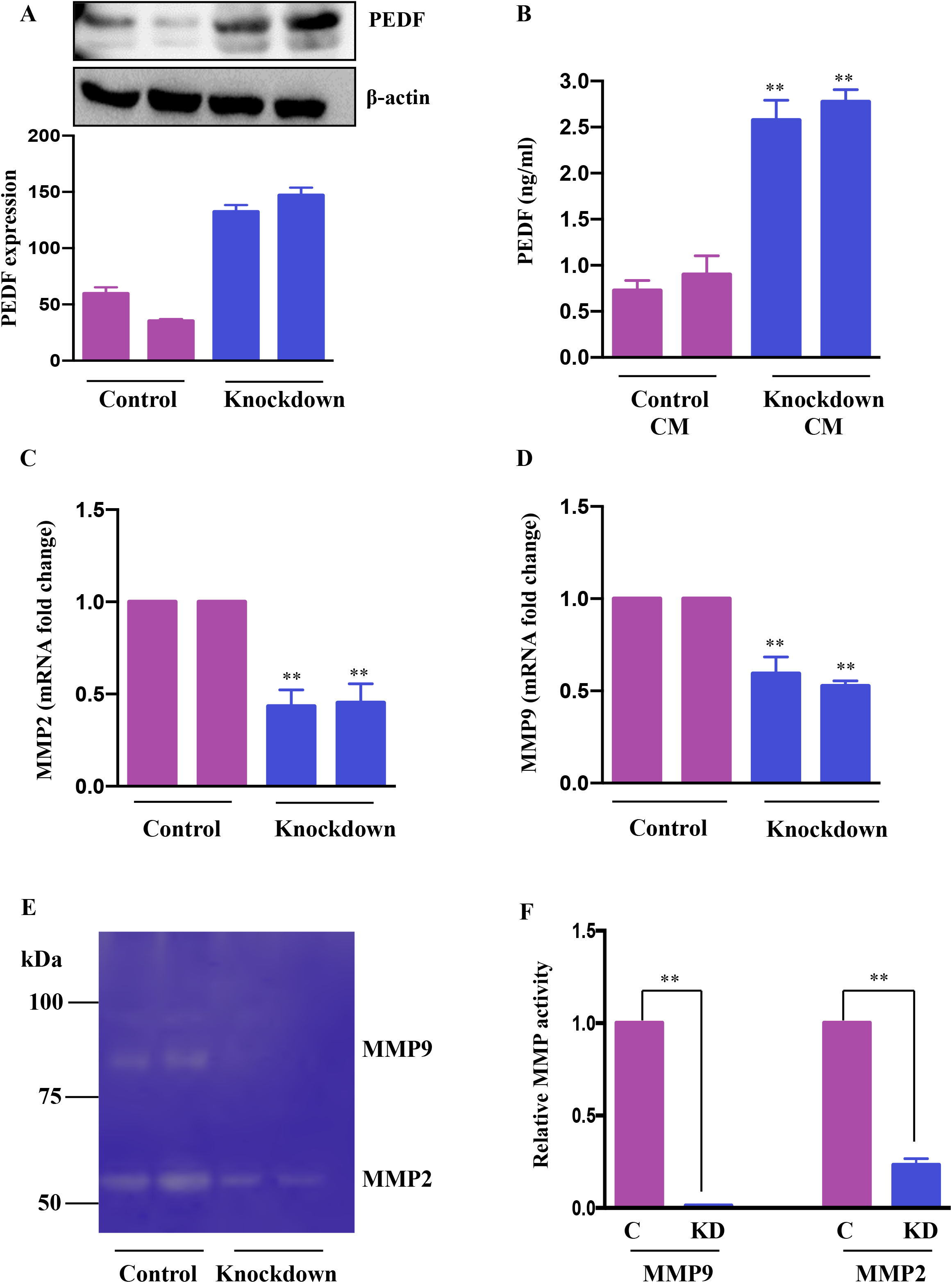
Knockdown of AATF increases PEDF expression. (A) The mRNA expression of PEDF in control and AATF knockdown QGY-7703 cells (B) PEDF levels in conditioned media (CM) from control and AATF knockdown cells was measured by ELISA. The mRNA expression of MMP2 (C) and MMP9 (D) in control and AATF knockdown QGY-7703 cells. (E) MMP2 and MMP9 gelatinolytic activity in the conditioned media of control and AATF knockdown cells was detected by gelatin zymography. (F) Quantitative densitometric analysis of MMP2 and MMP9 lysis bands of control and AATF knockdown QGY-7703 cells following zymography. Data are expressed as the mean ± SEM of three experiments. **p < 0.001 or *p < 0.05 compared to control.

We next explored the mechanism by which PEDF is upregulated in AATF knockdown cells. PEDF, a member of the serpin superfamily, acts as a natural angiogenesis inhibitor and is found to be significantly downregulated in most cancers, including HCC (33). There is an increasing body of evidence for the involvement of matrix metalloproteinases type 2 (MMP2) and type 9 (MMP9) in the degradation of PEDF. Of note, PEDF acts as a substrate for MMP2 and MMP9 (34). Along the same lines, we examined the expression of MMP2 and MMP9 in CM of control and knockdown QGY7703 cells and interestingly found that MMP2 and MMP9 were down regulated in AATF knockdown cells compared to control (**Figure 3C and 3D**). Additionally, we also tested the MMP activity by performing gelatin zymography. Consistent with the expression data, there was decreased gelatinolytic activity of MMP2 and MMP9 in knockdown cells compared to control (**Figure 3E and 3F**). Together, these findings suggested that AATF inhibition downregulates MMP2 and MMP9, which prevents PEDF from being degraded in AATF knockdown cells as opposed to control cells.

### AATF knockdown exerts anti-angiogenic effect in HCC via PEDF

Further investigations were carried out to determine whether PEDF played a key role in the suppression of angiogenesis in AATF knockdown cells. To test this hypothesis, we further validated the effect of conditioned media from control and AATF knockdown QGY-7703 cells on the migration and invasion of HUVECs and vascular formation in CAM in the presence of neutralizing anti-PEDF antibody. A non-specific IgG antibody served as an isotype control. The concentration of anti-PEDF antibody was determined by a dose response experiment on the proliferation of HUVECs with control and AATF knockdown CM, and a concentration of 5 μg/ml was chosen for the experiments. Consistent with the previous experiments, the conditioned media of AATF knockdown cells inhibited the migration (30% vs. 100% of cell migration in the control) and invasion (18 vs. 75 cells per field in the control) of HUVECs, and this inhibition was significantly diminished in the presence of PEDF neutralizing antibody [72% of cell migration (**Figure 4A and 4D**) and invasion assay-46 cells per field (**Figure 4B and E**)]. Similarly, the decrease in the vascular growth caused by the conditioned media of AATF knockdown cells (7 vs. 23 blood vessels in control) was diminished in the presence of PEDF neutralizing antibody (16 blood vessels) (**Figure 4C and 4F**). Taken together, these data provide solid evidence that knockdown of AATF suppresses angiogenesis in HCC via PEDF.

**Figure 4.**
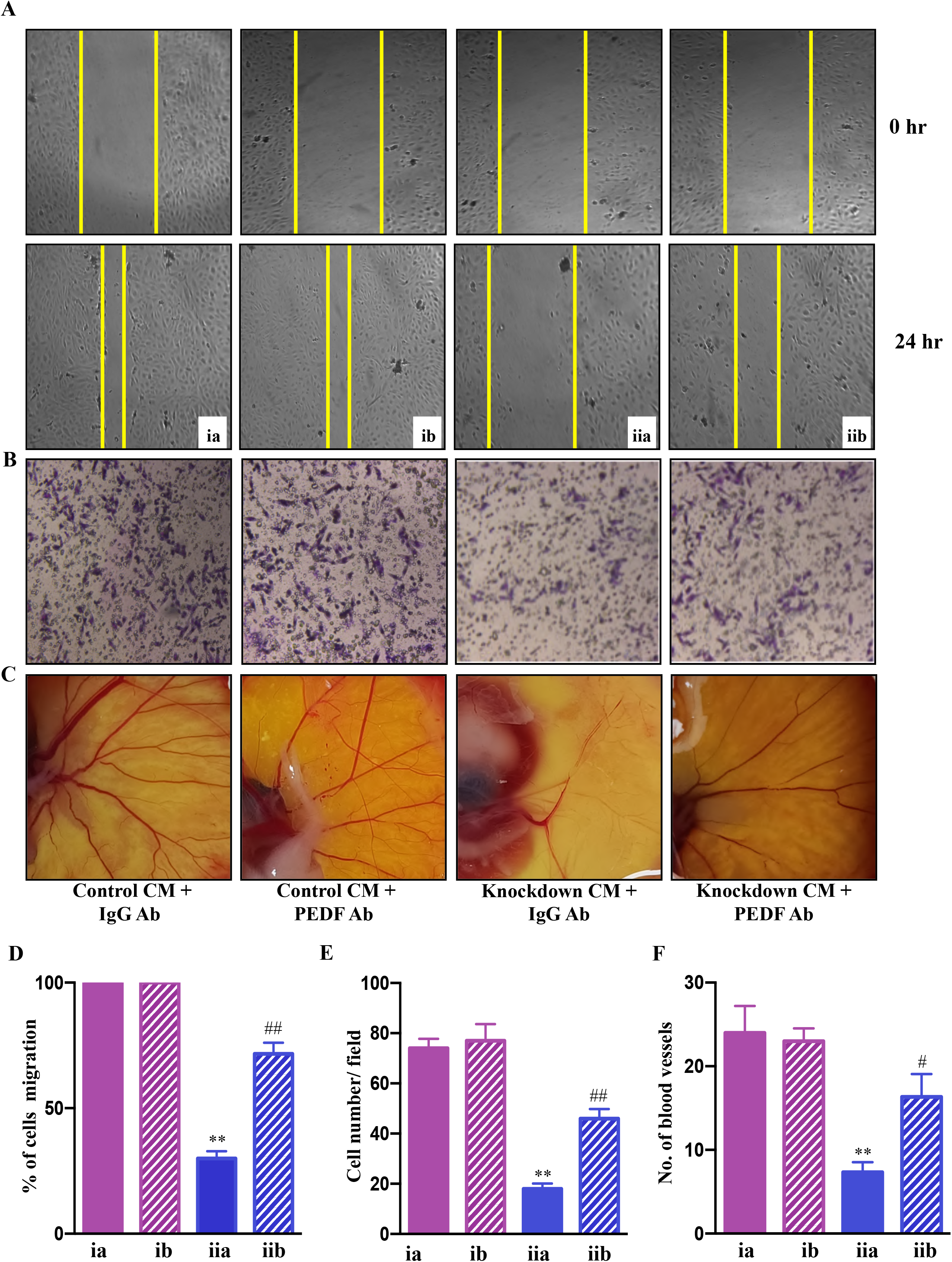
AATF knockdown inhibits angiogenesis via PEDF. Effect of conditioned media from control and AATF knockdown QGY-7703 cells treated with or without anti-PEDF antibody on migration (C) and invasion (D) of the human umbilical vein endothelial cells (HUVECs). (E) CAM assay performed using the conditioned media from control and AATF knockdown QGY-7703 cells treated with or without anti-PEDF antibody. Representative images are shown. (D) Quantification of the gap distance at 0 hr and 24 hr was evaluated using Image J software and expressed as % cells migration. (E) Quantitative analysis of HUVECs that passed through the membrane treated with conditioned media as characterized by matrigel invasion assay. (F) Quantification of the number of blood vessels was performed using Image J software. CM, conditioned medium; ia, control CM + IgG antibody; ib, control CM + PEDF antibody; iia, knockdown CM + IgG antibody; iib, knockdown CM + PEDF antibody. Data are expressed as the mean ± SEM of three experiments. **p < 0.001 or *p < 0.05 compared to ia; ##p < 0.001 or #p < 0.05 compared to iia.

### Mechanisms involved in AATF-mediated angiogenesis in human HCC

To examine the mechanisms underlying AATF-mediated angiogenesis in human HCC, we sought to explore several downstream signaling effectors that are responsible for endothelial cell survival and vascular permeability, cell proliferation, and migration. To test this hypothesis, HUVECs were treated with CM from control and AATF knockdown cells and examined for Erk1/2, Akt, and FAK phosphorylation by western blot analysis. Our results showed that the activation of the key components of the angiogenesis signaling pathway, including the phosphorylation of Erk1/2 (**Figure 5A**), Akt (**Figure 5B),** and FAK (**Figure 5C)**, was decreased in HUVECs treated with CM from knockdown cells compared to control. To corroborate the involvement of PEDF in the angiogenesis signaling pathway, experiments were carried out in the presence of neutralizing anti-PEDF antibody. Consistent with our previous observations, the activation i.e. phosphorylation of Erk1/2, Akt, and FAK, which was decreased by the AATF knockdown, was diminished in the presence of PEDF neutralizing antibody (**Figure 5A, 5B, and 5C**).

**Figure 5.**
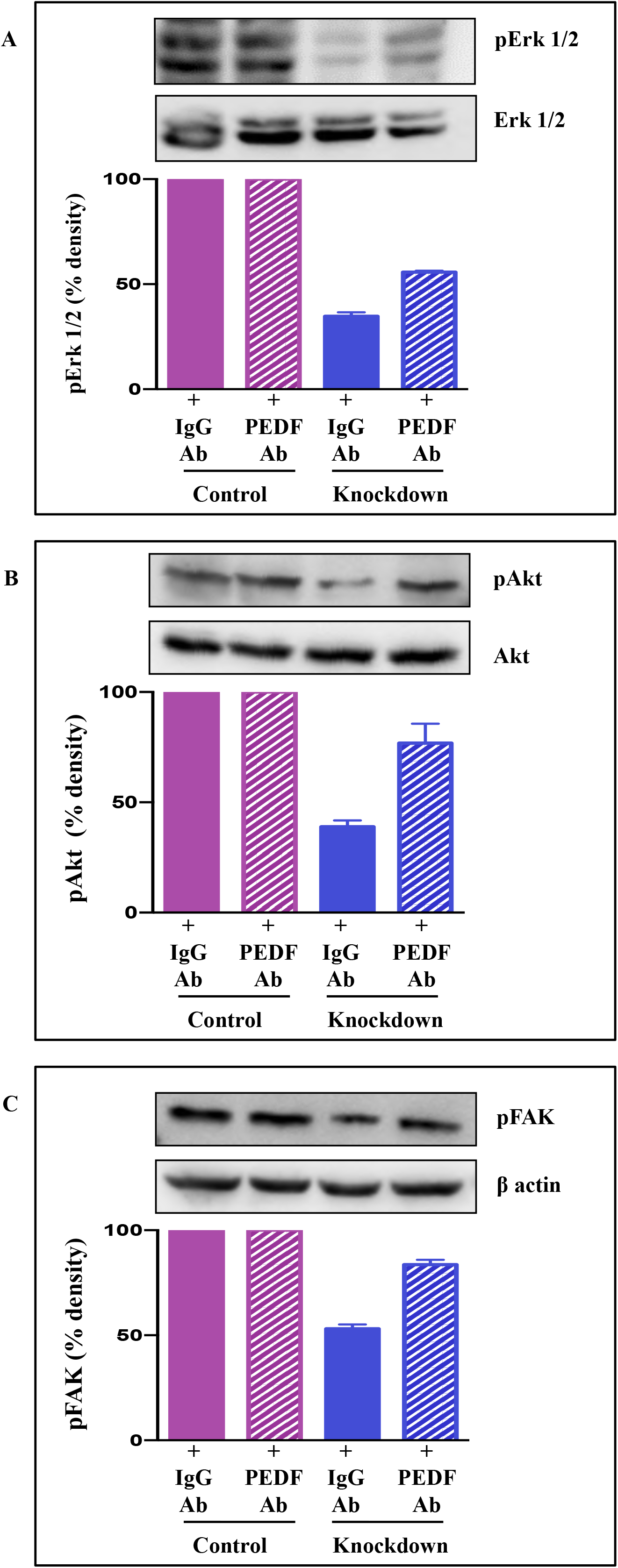
Signaling pathway via which AATF knockdown inhibits angiogenesis in HCC. Western blots depicting the protein expression of pErk1/2 and Erk1/2 (A), pAkt and Akt (B), and pFAK (C) in HUVECs treated with conditioned media from control and AATF knockdown QGY-7703 cells treated with or without anti-PEDF antibody. Bar graphs show the densitometric values calculated after normalization to their respective controls (pErk/Erk, pAkt/Akt and pFAK/β-actin). Data are expressed as the mean ± SEM of three experiments. **p < 0.001 or *p < 0.05 compared to control.

In our previous studies, we discovered that human HCC cells have elevated STAT3 levels as well as the AATF-STAT3 nuclear interaction (28). Studies by Zhang et al., have demonstrated that elevated activation of STAT3 is responsible for the upregulation of MMP2 and MMP9 in cancer cells (35). Taken together, it is evident that AATF interacts with STAT3 and upregulates the matrix metalloproteinases, MMP2 and MMP9, which in turn degrade PEDF. In contrast, inhibiting AATF reduces MMP2 and MMP9 levels, effectively stopping the degradation process, while PEDF remains functionally active. Of note, it is known that PEDF inhibits VEGF-induced phosphorylation and activation of VEGFR (36). Thus, we provide experimental evidence of how inhibition of AATF exerts an anti-angiogenic effect in HCC via PEDF. This results in decreased cell proliferation, cell survival, vascular permeability, and cell migration, which in turn leads to the inhibition of angiogenesis, preventing tumor growth in HCC (**Figure 6**). All these data demonstrated that AATF inhibition provided positive effects on human HCC.

**Figure 6.**
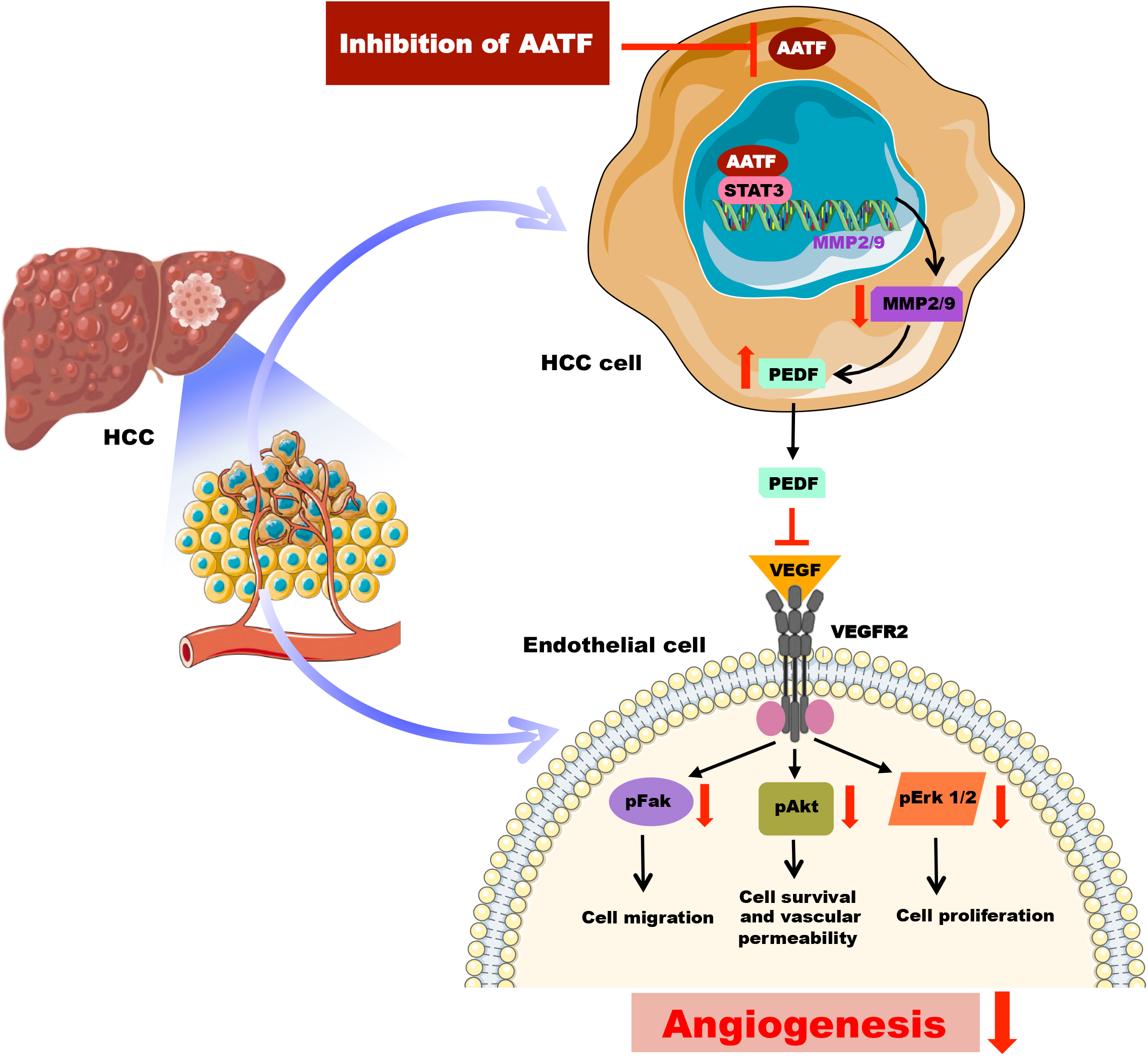
Schematic representation of the molecular mechanisms involved in AATF-mediated regulation of angiogenesis in human HCC.

## DISCUSSION

The current manuscript describes highly significant and relevant findings in the context of HCC, a highly vascularized tumor in which angiogenesis is critical to its growth, invasion, and metastasis (31). While AATF expression is very low or non-existent in normal liver tissues, it gradually increases as the stages and grades of HCC disease progress. Numerous studies have discovered that AATF, a multifunctional and highly conserved protein, contributes significantly to the development of various malignancies (14,17,21). We have previously shown the novel regulatory role of AATF in NAFLD-associated HCC (28). This is the first study to elucidate that targeting AATF can be an effective antiangiogenic strategy in the treatment of HCC.

Our findings in NAFLD-associated HCC showed that AATF expression was upregulated, whereas knocking down AATF significantly reduced tumor burden and metastasis in a mouse xenograft model (28). These data led to investigations to understand the role of AATF in the molecular pathogenesis of HCC, and therefore we elucidated the unexplored regulatory role of AATF in tumor angiogenesis, one of the hallmarks that contribute to tumor growth and metastasis. To precisely elucidate the molecular mechanism underlying AATF function in angiogenesis, we focused on knocking down AATF in QGY-7703 human HCC cells. The first finding is that the CM of AATF knockdown cells inhibited the proliferation of human umbilical vein endothelial cells (HUVECs) compared to the control. The data are concordant with the migration and invasion of HUVECs, wherein AATF knockdown caused inhibition compared to control. We also document that the vascular growth in chicken embryos is inhibited with CM of AATF knockdown HCC cells, as assessed by the CAM assay, providing strong experimental evidence that inhibition of AATF suppresses angiogenesis in HCC.

Tumor-induced angiogenesis is typically associated with a complex interplay of multiple factors and pathways, with vascular endothelial growth factor (VEGF) being a key player (37). The process of angiogenesis in HCC is an extremely complex and tightly regulated process characterized by well-balanced interactions between pro- and anti-angiogenic factors. In addition to VEGF, the other key angiogenesis stimulating factors include PEDF, FGF, angiopoietins, and endoglins. On the other hand, endogenous angiogenesis inhibitors include anti-angiogenic peptides, hormone metabolites, and apoptosis modulators (8,38). Thus, the angiogenic switch involving the proangiogenic factors overexpression as well as anti-angiogenic factors inhibition results in increased tumor vascular burden, which leads to tumor proliferation and progression (39). Perhaps, the strongest evidence to support the role of AATF in angiogenesis came from our previous studies which showed the increase of PEDF or SerpinF1 levels in the AATF knockdown HCC cells compared to control as demonstrated by the human angiogenesis array (28). This is further corroborated in the current study, which confirmed high levels of PEDF, both by immunoblot analysis and ELISA, in AATF knockdown cells compared to control cells. Similarly, Matsumato K. et al., found lower PEDF serum concentrations in patients with cirrhosis or HCC compared to healthy subjects (40). This provided a strong rationale to further examine the involvement of PEDF in AATF-mediated angiogenesis in HCC.

PEDF, belonging to the serine protease inhibitor (serpin) superfamily, has several roles that frequently work against the pathways that promote the progression of cancer (41). Notably, Dawson and his colleagues for the first time identified the potent anti-angiogenic activity of PEDF in the cornea and vitreous humour of the eye (42), which later paved the way for the exploration of PEDF as an angiogenesis inhibitor in tumors (43). PEDF, on the other hand, is a substrate for extracellular matrix metalloproteinases and has been found to be degraded in several cancers (34,41). In this regard, our findings revealed a decrease in MMP2 and MMP9 activity in the CM of AATF knockdown cells compared to controls, providing a logical explanation for the presence of PEDF levels in AATF knockdown HCC cells versus controls. These data are in support of the studies carried out by Notari L. et al., which showed MMP-mediated degradation of PEDF leading to increased angiogenesis in the retina (34). Furthermore, the impact of PEDF in AATF-mediated angiogenesis was supported by the observation that the inhibition of PEDF by anti-PEDF antibody significantly diminished the inhibition of migration and invasion of HUVECs and inhibition of vascular growth in chicken embryos treated with CM of AATF knockdown cells. It is interesting to note that, PEDF inhibits angiogenesis either by increasing γ-secretase-mediated cleavage of VEGFR2 or by inhibiting VEGF-induced phosphorylation and activation of VEGFR2 (44). However, in our studies, the activation of Erk1/2, Akt, and FAK, which are the angiogenic mediators, downstream of VEGF signaling, was inhibited in HUVECs treated with CM of knockdown cells. Also, this inhibition was diminished in the presence of the PEDF antibody. These data are thus in line with the mechanism of PEDF antagonizing the action of VEGF-A binding to its receptor, VEGFR2 and thereby preventing the activation and downstream VEGF-A signals (36). From the mechanistic point of view, our current study provides an insight into the mechanisms of AATF inhibition exerting anti-angiogenic effects in human HCC via PEDF (**Figure 6**). This also offers an additional advantage, as PEDF not only blocks angiogenesis but is also involved in anti-tumor and anti-metastatic activities.

This proof-of-concept study supporting the notion that AATF inhibition suppresses angiogenesis holds certain clinical implications. Clinically, liver-specific targeting of AATF by lipid nanoparticle-based delivery of siRNA-AATF or AAV8-mediated siRNA-AATF delivery appears to be an effective antiangiogenic approach. Another approach would be the generation of liver-specific AATF knockout mice that can be used to induce HCC and examine the role of AATF in tumor progression, angiogenesis, and metastasis. Though the FDA-approved angiogenesis inhibitors such as sorafenib, bevacizumab, and ramucirumab possess efficacy, they are known to have adverse effects, and resistance is a major concern (45). There also exists the therapeutic limitation of systemic administration of antiangiogenic compounds restricting their clinical applications (46). Of note, this antiangiogenic gene therapy involving AATF knockdown may be an attractive strategy due to its specificity. Unlike the angiogenic inhibitors/compounds that may inhibit growth factor-induced signal transduction that is required in tumor angiogenesis as well as normal vasculature, resulting in adverse effects. Thus, targeting angiogenesis by AATF inhibition would be a safe antiangiogenic approach. This also warrants future detailed preclinical studies involving AATF inhibition and understanding the molecular mechanisms of AATF-mediated metastasis in HCC. It is of prime importance that current strategies for combating angiogenesis and the potential for combining antiangiogenic therapy with other systemic modalities, like immunotherapy, would help HCC patients have a better prognosis.

In summary, the current study adds to the growing body of evidence supporting the key role of angiogenesis in tumor growth and progression. We demonstrated that AATF inhibition suppresses angiogenesis in human HCC via PEDF, and AATF may serve as a promising gene therapy for HCC treatment. The study offers a target for intervention and a direction for investigations to lessen the burden of HCC by targeting angiogenesis, and it provides a basis for future work that can be translated into human trials.

## ACKNOWLEDGMENT

We gratefully acknowledge Dr. Suma M.N., Vice Principal and Head of the Department of Biochemistry, JSS Medical College, for her guidance, as well as Dr. Nalini and her team, Gynecologist, Shree Devi Nursing Home, Mysore, for providing human umbilical cord for the study. Human normal or HCC tissues were obtained from the National Tumor Tissue Repository (NTTR), Tata Memorial Hospital, Mumbai.

## AUTHOR CONTRIBUTIONS

DS designed and performed experiments, analyzed data, and wrote the manuscript; ANS performed experiments and analyzed data; SS analyzed the data; AP and PV contributed to the discussion and reviewed the manuscript; PKS provided scientific insights and reviewed the manuscript; and DPK conceptualized the project, designed experiments, critically evaluated the results, and wrote the manuscript. All authors contributed to the article and approved the final version of the manuscript submitted.

## ABBREVIATIONS

HCC: hepatocellular carcinoma
AATF: apoptosis antagonizing transcription factor
DNA: deoxyribonucleic acid
UPR: unfolded protein response
ER: endoplasmic reticulum
NAFLD: nonalcoholic fatty liver disease
HUVECs: human umbilical vein endothelial cells
VEGF: vascular endothelial growth factor
PDGF: platelet-derived growth factor
FGF: fibroblast growth factor
PEDF: pigment epithelium-derived factor
CAM: chorioallantoic membrane
CM: conditioned media
nrf2: nuclear factor erythroid-related factor 2
NRAGE: neurotrophin receptor-interacting MAGE homolog
MMP: matrix metalloproteinases
ERK1/2: extracellular signal-regulated protein kinase
FAK: focal adhesion kinase
STAT3: signal transducer and activator of transcription 3
VEGFR: vascular endothelial growth factor receptor
siRNA: small interfering ribonucleic acid
AAV: adeno associated virus

---

**Supplementary figure 1.** Culturing of Human umbilical vein endothelial cells (HUVECs). Images at different magnifications-10X, 20X and 40X.

**Supplementary figure 2.** Analysis of AATF expression in normal (n=50) and human HCC (n=371) tissues in TCGA microarray data set (A). Expression of AATF varies across the stages of HCC (B) and tumor grade (C).

**Supplementary figure 3.** AATF mRNA expression in control and AATF knockdown QGY-7703 cells. Data are expressed as the mean ± SEM of three experiments. **p < 0.001 or *p < 0.05 compared to control.

**Supplementary Table 1.** Demographic and clinicopathological data of HCC patients. M, male; F, female; HCC, hepatocellular carcinoma; TNM, tumor, nodes and metastases; PD, poorly differentiated; MD, moderately differentiated; WD, well differentiated.

**Supplementary Table 2.** List of primers used in qRT-PCR

