## Supplementary material for "AATF Inhibition Exerts Antiangiogenic Effects Against Human Hepatocellular Carcinoma"

**Supplementary Figure 1**

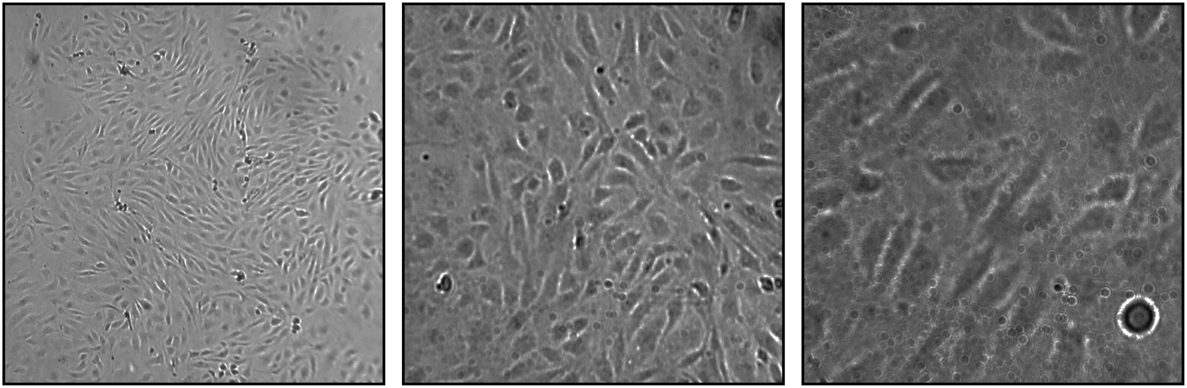

10X 20X 40X

**Supplementary Figure 2**

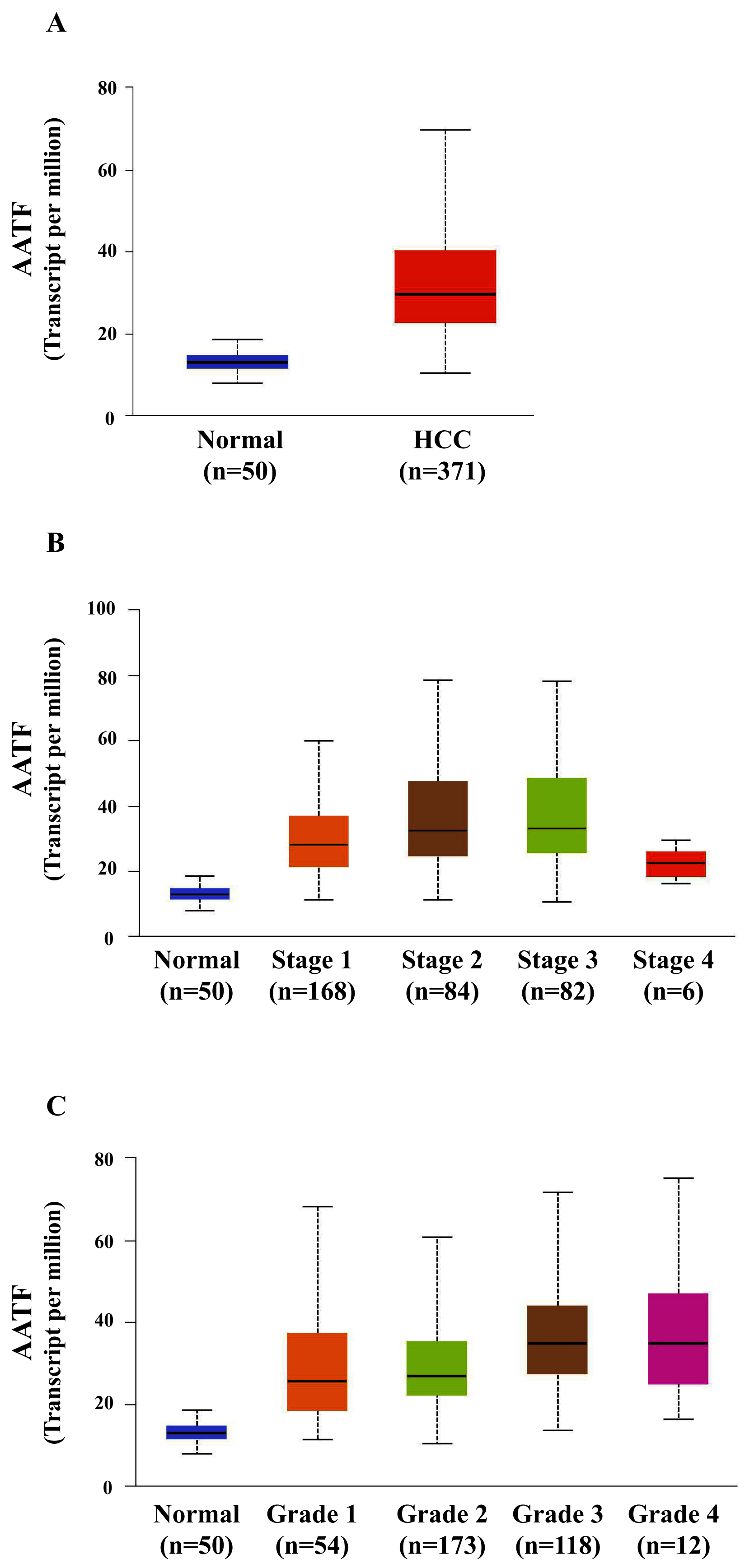

**Supplementary Figure 3**

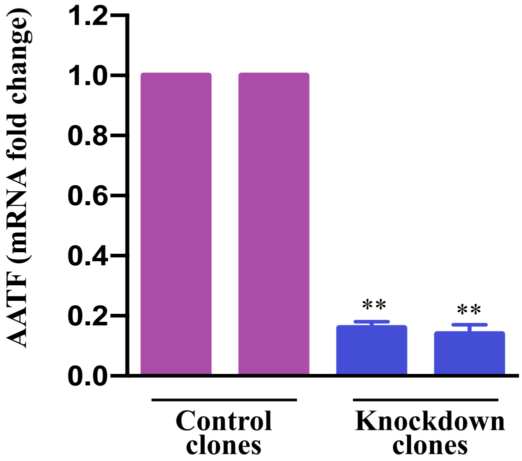

**Supplementary Table 1**

| Sl. No. | Gender | Age (yr) | Patient Diagnosis | TNM | Tumor Stage | Tumor Grade |
|---|---|---|---|---|---|---|
| 1. | F | 38 | HCC | pT1bNxM0 | Stage 1 | PD |
| 2. | M | 78 | HCC | pT1bN0M0 | Stage 1 | MD |
| 3. | M | 64 | HCC | pT1bNxM0 | Stage 1 | MD |
| 4. | F | 35 | HCC, infiltrating stomach | pT4NxM0 | Stage 4 | MD |
| 5. | F | 56 | HCC of left lobe of liver | pT3N1M0 | Stage 3 | MD |
| 6. | M | 81 | HCC | pT3N0M0 | Stage 3 | MD |
| 7. | M | 43 | HCC | pT1bNxM0 | Stage 1 | MD |
| 8. | F | 72 | HCC | pT2N0M0 | Stage 2 | MD |
| 9. | M | 55 | HCC | pT1bNxM0 | Stage 1 | MD |
| 10. | F | 49 | HCC | pT1bNxM0 | Stage 1 | MD |
| 11. | M | 85 | HCC | pT1bNxM0 | Stage 1 | MD |
| 12. | F | 30 | HCC | pT3aN0MX | Stage 3 | MD |
| 13. | F | 70 | HCC | pT3NxM0 | Stage 3 | PD |
| 14. | F | 67 | HCC | pT3N0M0 | Stage 3 | WD |
| 15. | M | 72 | HCC | pT3NxM0 | Stage 3 | PD |
| 16. | M | 42 | HCC | pT3N0M0 | Stage 3 | PD |
| 17. | M | 46 | HCC | pT2NxM0 | Stage 2 | PD |
| 18. | M | 81 | HCC | pT3NxM0 | Stage 3 | MD |
| 19. | M | 65 | HCC | pT1bN0M0 | Stage 1 | MD |
| 20. | M | 69 | HCC | pT3N0M0 | Stage 3 | MD |
| 21. | M | 44 | HCC | pT2NxM0 | Stage 2 | PD |

| | | | | | | |
|---|---|---|---|---|---|---|
| 22. | M | 67 | HCC | pT2NxM0 | Stage 2 | MD |
| 23. | M | 51 | HCC | pT4N0 | Stage 4 | MD |
| 24. | M | 35 | HCC | pT2NxM0 | Stage 2 | MD |
| 25. | M | 66 | HCC | pT1NxMx | Stage 1 | MD |
| 26. | M | 59 | HCC | pT4NxM0 | Stage 4 | PD |
| 27. | M | 68 | HCC, anaplastic type | pT3NxM0 | Stage 3 | PD |
| 28. | M | 51 | HCC | pT2NxM0 | Stage 2 | MD |
| 29. | M | 67 | HCC | pT2N0M0 | Stage 2 | MD |
| 30. | M | 46 | HCC | pT1bN0M0 | Stage 1 | MD |
| 31. | M | 73 | HCC | pT2NxM0 | Stage 2 | MD |
| 32. | F | 52 | HCC | pT3NxM0 | Stage 3 | PD |
| 33. | M | 80 | HCC | pT1bNxM0 | Stage 1 | MD |
| 34. | M | 69 | HCC | pT3N0M0 | Stage 3 | MD |
| 35. | M | 54 | HCC | pT1bNxM0 | Stage 1 | PD |
| 36. | F | 44 | HCC | pT2NxM0 | Stage 2 | MD |
| 37. | M | 53 | HCC | pT2NxM0 | Stage 2 | PD |
| 38. | F | 61 | HCC | pT3N0M0 | Stage 3 | MD |
| 39. | M | 77 | HCC | pT2 | Stage 2 | PD |
| 40. | M | 71 | HCC with predominant undifferentiated morphology | pT3NxM0 | Stage 3 | PD |
| 41. | M | 71 | HCC | T3N0M0 | Stage 3 | PD |
| 42. | F | 54 | HCC | pT1bN0M0 | Stage 1 | MD |
| 43. | M | 49 | HCC | pT3NxM0 | Stage 3 | MD |

| | | | | | | |
|---|---|---|---|---|---|---|
| 44. | M | 75 | HCC | pT2NxM0 | Stage 2 | PD |
| 45. | M | 25 | HCC, fibrolamellar | pT3NxM0 | Stage 3 | PD |
| 46. | M | 51 | HCC | pT2N0 | Stage 2 | WD |
| 47. | M | 54 | HCC | pT1NxM0 | Stage 1 | WD |
| 48. | M | 72 | HCC | pT3NXMX | Stage 3 | MD |
| 49. | F | 36 | HCC | pT1bN0M0 | Stage 1 | MD |
| 50. | M | 45 | HCC | pT2pN0pM0 | Stage 2 | MD |

**Supplementary Table 2**

| Name | Forward Primer | Backward Primer |
|---|---|---|
| **AATF** | TAGAACGGAAGACCAGCTCC | TGCTAAGGACATGAAACCGA |
| **β- actin** | AGAGATGGCCACGGCTGCTT | CAGGACTCCATGCCCAGGAA |
| **PEDF** | GGTGCAGGCGCAGATGAAAG | TTGTTACCCACTGCCCCTTGA |
| **MMP2** | TACGATGGAGGCGCTAATGGC | GAAGGTGTTCAGGTATTGCACTG |
| **MMP9** | TCTGCCCGGACCAAGGATAC | CCCCTCAGTGAAGCGGTACAT |
